# Advanced computational and statistical multiparametric analysis of Susceptibility-Weighted Imaging to characterize gliomas and brain metastases

**DOI:** 10.1101/2020.04.24.060830

**Authors:** Antonio Di Ieva, Carlo Russo, Pierre-Jean Le Reste, John S. Magnussen, Gillian Heller

## Abstract

Susceptibility-weighted imaging (SWI) is a technique useful for evaluation of the internal structures of brain tumors, including microvasculature and microbleeds. Intratumoral patterns of magnetic susceptibility can be quantified by means of fractal analysis. Here, we propose a radiomics methodological pipeline to merge advanced fractal-based computational modelling with statistical analysis to objectively characterize the fingerprint of gliomas and brain metastases. Forty-seven patients with glioma (grades II-IV, according to the WHO 2016 classification system) and fourteen with brain metastases underwent 3 Tesla MRI using a SWI protocol. All images underwent computational analysis aimed to quantify three Euclidean parameters (related to tumor and SWI volume) and five fractal-based parameters (related to the pixel distribution and geometrical complexity of the SWI patterns). Principal components analysis, linear and quadratic discriminant analysis, K-nearest neighbor and support vector machine methods were used to discriminate between tumor types. The combination of parameters offered an objective evaluation of the SWI pattern in gliomas and brain metastases. The model accurately predicted 88% of glioblastoma, according to the quantification of intratumoral SWI features, failing to discriminate the other types. SWI is not normally used to classify brain tumors, however fractal-based multi-parametric computational analysis can be used to characterize intratumoral SWI patterns to objectively quantify tumors-related features. Specific parameters still have to be identified to provide completely automatic computerized differential diagnosis.

## 1 Introduction

With the introductions of more complex and potentially more sensitive imaging techniques in neuroradiology, there is a growing need for quantitative analysis to provide objective measures of radiological features in conjunction with the qualitative description by the specialist radiologist. Artificial intelligence techniques are being rapidly deployed in neuroradiology and radiomics [1, 2], many of which may benefit from new image markers to describe and characterize regions of interest (ROI), quite apart from new computerized parameters which could quantify specific features in an objective way.

Susceptibility-weighted imaging (SWI) is an imaging technique useful for the evaluation of the internal structures of brain tumors, including microvasculature and microbleeds [3, 4]. SWI can demonstrate intra-tumor heterogeneity; that is, the intralesional signal patterns and edge irregularity[5].

In the field of quantitative neuroradiology and clinical neurosciences, fractal-based computational modelling has been introduced for the characterization of intrinsic features (e.g., microvascular patterns) [6–8]. The intratumoral patterns of magnetic susceptibility can be quantified by means of fractal analysis [9]. In prior research, we have demonstrated that the fractal dimension of intratumoral SWI patterns, as seen at ultra-high-field (7 Tesla) [5] and high-field (3 Tesla) [10], is a valid morphometric image marker to estimate glial brain tumor grade, which is useful in cancer and neuroimaging research as well as in the clinical setting.

Here, we propose a methodological pipeline merging advanced fractal-based computational modelling with statistical analysis to objectively characterize the fingerprint of gliomas and brain metastases. The current model aims to generate neuroimaging fingerprints of brain tumors, to help neuroradiologists and clinicians to characterize and follow-up patients undergoing serial radiological investigations.

## 2 Materials and methods

### 2.1 Patients

Following informed consent and local ethics committee approval, we retrospectively analyzed the pre-surgical imaging of 61 patients (mean age 51 +/- 16 years; range 24-77) who underwent multimodal imaging, including SWI. The pre-operative images were compared with the post-surgical pathological analysis of the sample, consisting of: 13 grade II gliomas (9 oligodendroglioma, 1 oligoastrocytoma, 3 astrocytoma); 10 grade III gliomas (7 anaplastic oligodendrogliomas, 3 anaplastic astrocytomas), and 24 grade IV gliomas (glioblastoma); 14 metastases (3 melanomas, 3 mammary carcinoma, 6 pulmonary adenocarcinomas, 1 gallbladder carcinoma, and 1 uterine carcinoma).

### 2.2 MR Imaging

All MRIs were performed on a 3-T Philips Achieva scanner (Philips, Amsterdam, the Netherlands). The sequences used in this study included T1-weighted isotropic gradient-echo imaging before and after gadolinium infusion, T2 fluid-attenuated inversion recovery imaging, and 2 types of 3-dimensional SWI depending on the year of acquisition (matrix = 512 x 512, resolution = 0.43 x 0.43 x 0.5 mm, 200 slices, repetition time/echo time = 22/32 ms or matrix = 512 x 512, resolution = 0.43 x 0.43 x 1.2mm, 84 slices, repetition time/echo time = 28.3/38.3 ms). The tumors were contoured by consensus on each slice of native susceptibility-weighted images by a neurosurgeon and a neuroradiologist who were blinded to the histology results. The segmentation was produced with the help of all sequences rigidly registered on SWI using a 12-degree of freedom algorithm (MIPAV - Medical Image Processing, Analysis, and Visualization, which is freely distributed at http://mipav.cit.nih.gov/).

### 2.3 Postprocessing image analysis

All the images were filtered by using the Brightness Progressive Normalization algorithm (http://www.fractal-lab.org/Downloads/bpn_algorithm.html), designed to homogenize the grayscale level of the SWI signals across the dataset [9]. Following segmentation, each region of interest (ROI) represented the tumor volume. Three Euclidean parameters were calculated from the ROI: volume (defined as P5, see ***Table 1***); the threshold volume (defined as the volume of the structure obtained by intersecting the ROI and the gray level threshold) (P6); and the ratio between the threshold volume and the ROI volume (P7).

**Table 1.**
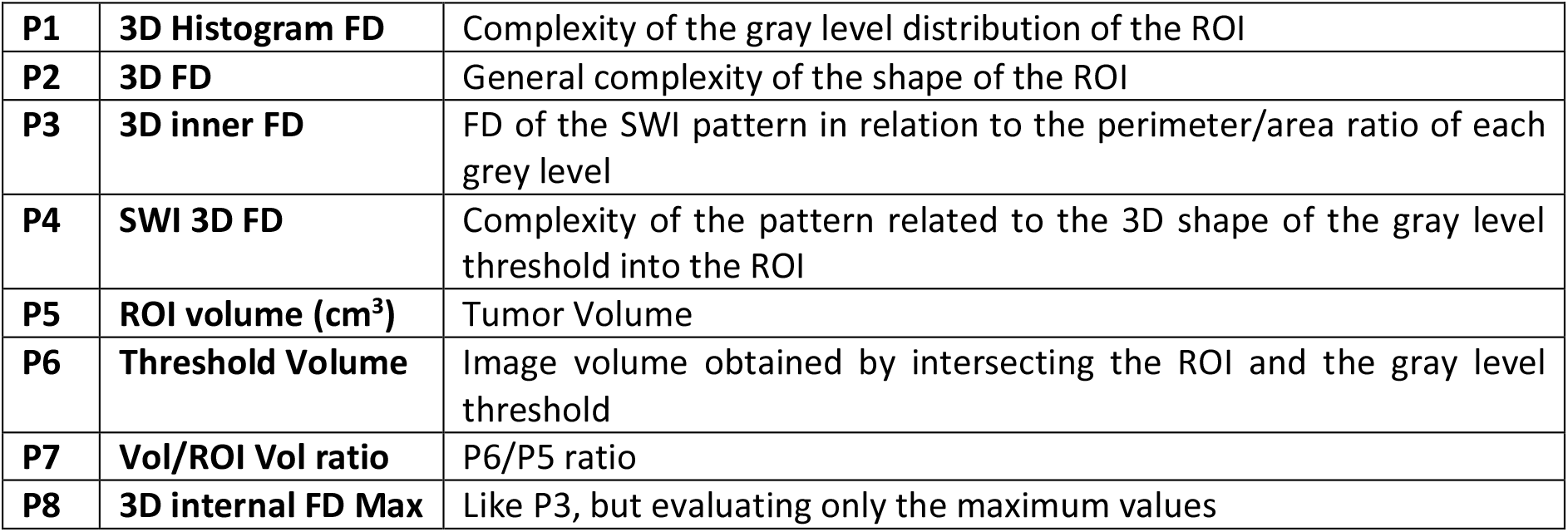
Definitions of the computed Euclidean (P5-P7) and fractal-based (P1-P4 and P8) parameters. FD: fractal dimension.

|  |  |  |
| --- | --- | --- |
| **P1** | **3D Histogram FD** | Complexity of the gray level distribution of the ROI |
| **P2** | **3D FD** | General complexity of the shape of the ROI |
| **P3** | **3D inner FD** | FD of the SWI pattern in relation to the perimeter/area ratio of each grey level |
| **P4** | **SWI 3D FD** | Complexity of the pattern related to the 3D shape of the gray level threshold into the ROI |
| **P5** | **ROI volume (cm<sup>3</sup>)** | Tumor Volume |
| **P6** | **Threshold Volume** | Image volume obtained by intersecting the ROI and the gray level threshold |
| **P7** | **Vol/ROI Vol ratio** | P6/P5 ratio |
| **P8** | **3D internal FD Max** | Like P3, but evaluating only the maximum values |

### 2.4 Fractal analysis

The box counting method was applied to assess the fractal dimension (FD). We used custom software developed in C++ for image and fractal analysis (http://www.fractal-lab.org/Downloads/FDEstimator.html). After image normalization and binarization, 5 fractal parameters were evaluated (**Fig. 1**), based on the FD, which is a non-integer number between 0 and 3 quantifying the geometric complexity of the analyzed object [11, 12]:

1. the 3D Histogram fractal dimension (P1), defined as the FD computed on the histogram of gray values in the selected ROI, evaluating gray level distribution rather than pixel arrangement. The value is obtained considering the distribution of the frequencies of each gray level of the pixels in the selected volume. The complexity of this distribution is given by the FD obtained evaluating the means of subsets of the distribution on different scales. This parameter evaluates gray level distribution but not the arrangement of the pixels, i.e. the same pixels with a different arrangement will have the same 3D Histogram FD.
2. the 3D FD (P2), defined as the FD of the 3D surface of the ROI shape. The ROI selected on each image forms a 3D object, the complexity of which is variable. This complexity is evaluated by means of the 3D box counting algorithm [11]. This parameter is able to evaluate complexity of the ROI shape but does not consider the pixel information of the images.
3. the 3D inner FD (P3), defined as the FD of the 3D internal grey levels distribution, obtained by the perimeter/area ratio of each contour line obtained sliding the gray level threshold; each grey level generates a shape, and the complexity of the non-linear perimeter/area ratio graph is evaluated by its fractal dimension. In this way all the useful information of the gray level values of the pixels and their distribution are considered in the FD evaluation. It is a more sensitive parameter compared to the SWI 3D FD (see below) and the 3D Histogram FD. This parameter is related to the change of the pixel gray value, its position, or both. It considers a more general rather than local complexity.
4. the SWI 3D FD (P4), defined as the FD of the 3D surface of the selected structure obtained by intersecting the ROI with a gray level threshold of the pixels, i.e., pixels that have the gray level included into a selected threshold of gray values and are inside the selected ROI. This parameter evaluates the general complexity of the SWI patterns. The segmentation of the SWI pattern forms a 3D object with a more or less complex 3D shape. The measure of the complexity of this shape is obtained by evaluating its FD using the 3D box counting algorithm. This parameter is able to evaluate the complexity of the shape but does not change when the internal gray levels of the pixel are different, when all the gray levels remains included into the selected threshold.
5. the 3D internal FD max (P8), defined as the 3D internal FD (like P3), but evaluating the FD on a log-log plot having only values up to the maximum extension of the perimeter in the perimeter/area ratio, making it an estimator of the distribution of the gray level and pixel distribution.

**Fig. 1.**
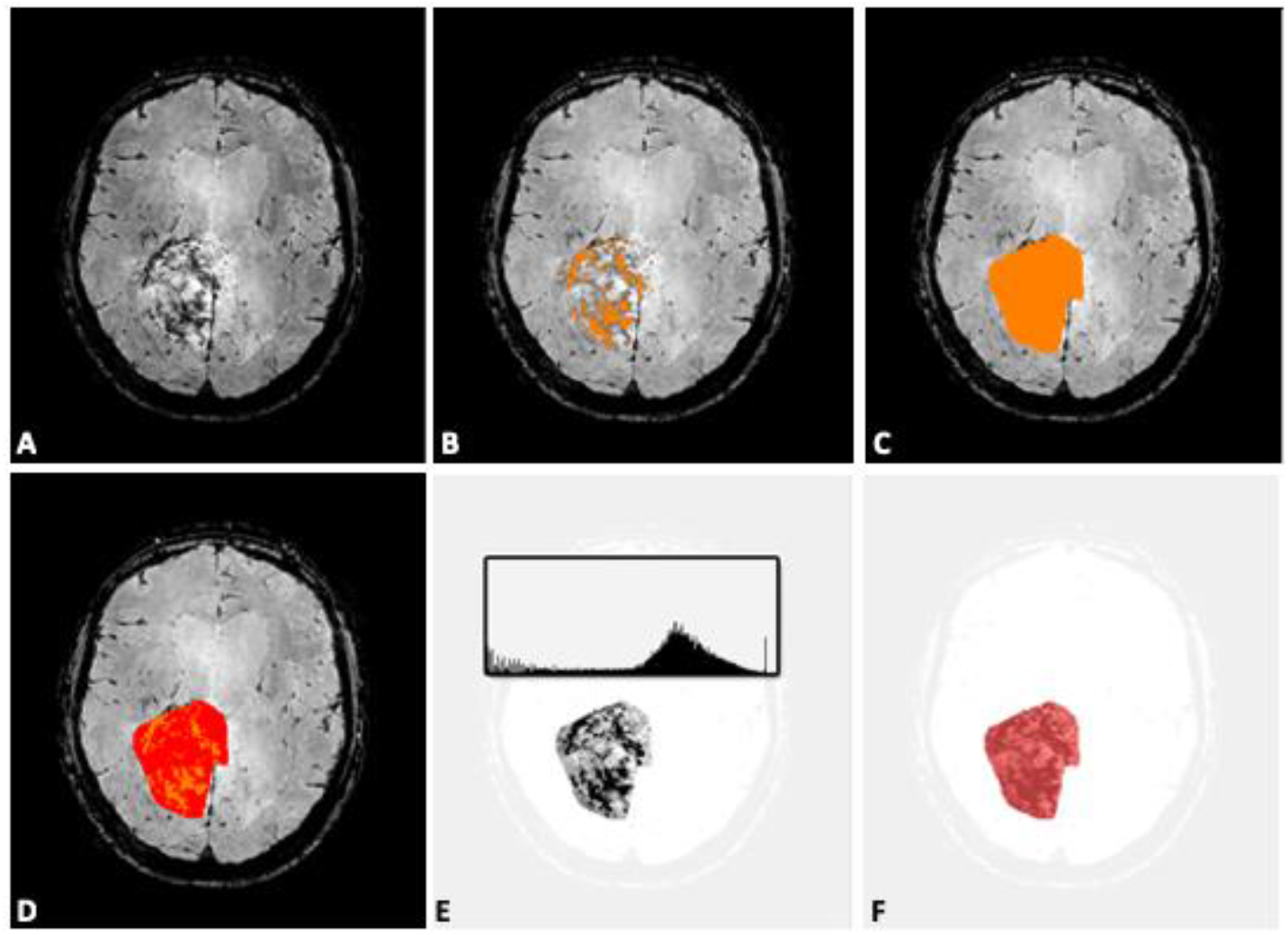
A) SWI of a right-sided glioblastoma. B) Automatic segmentation of the intratumoral SWI pattern for computing of the 3D FD (P2), evaluating the complexity of the heterogeneity of the SWI signal. C) ROI volume (P5) on the entire slices’ stack. D) Volume/ROI volume ratio. E) 3D Histogram FD (P1), evaluating the gray levels distribution; F) 3D inner FD (P3), evaluating the gray level and pixels’ distribution.

### 2.5 Statistical analysis

The data was pre-processed using Box-Cox transformations to normality [13], in order to maximize discrimination between groups. Principal component analysis was performed on the transformed variables, and the principal components explaining most of the variation were carried forward to the discrimination procedures. Linear and quadratic discriminant analysis, K-nearest neighbor and support vector machine methods [14] were implemented in order to assess their ability to discriminate between types.

## 3 Results

A matrix of plots of the raw data is given in ***Fig. 2***.

**Figure 2.**
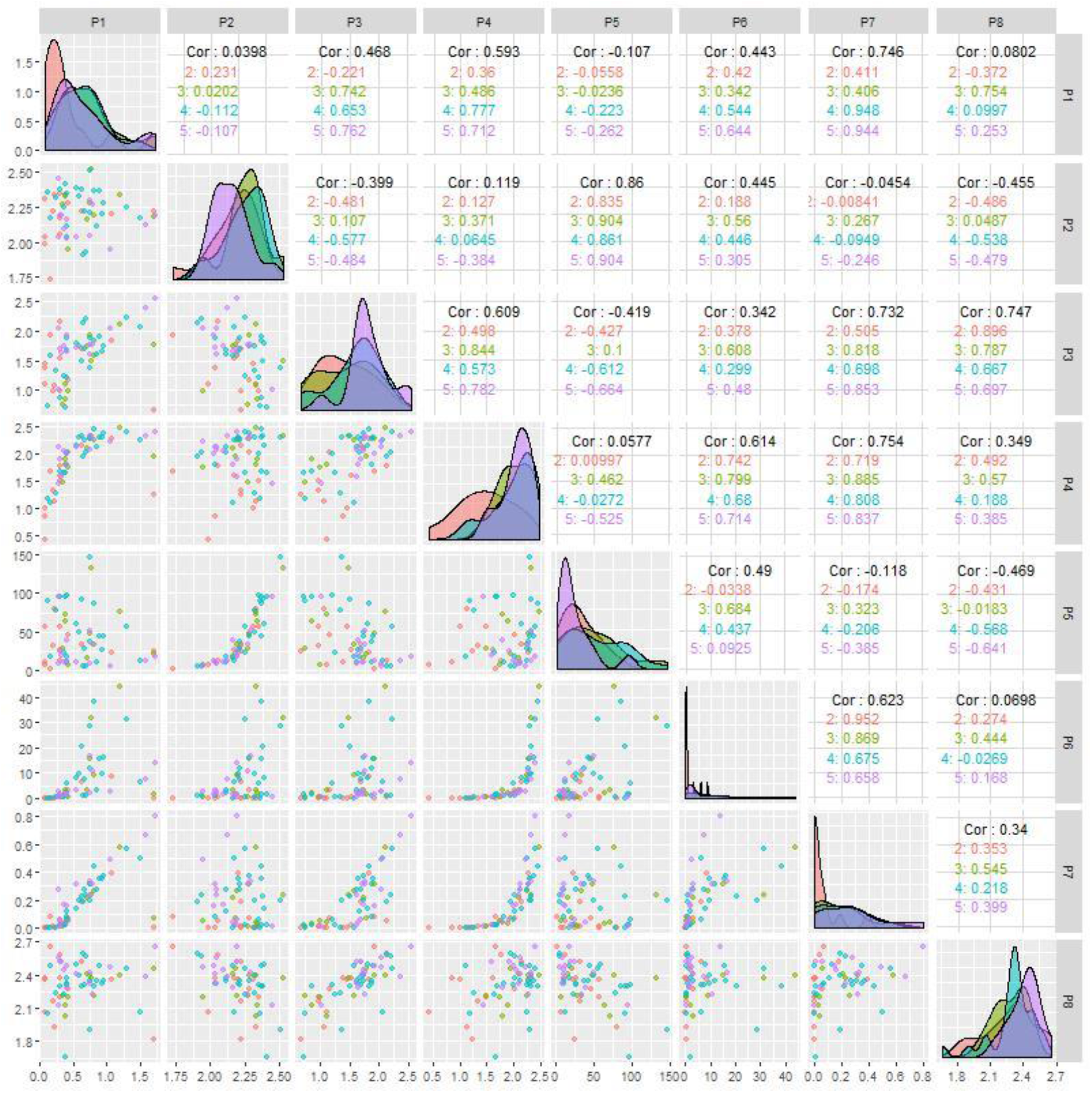
Matrix of plots of raw data. Density estimates by type are shown on the diagonal; bivariate scatterplots between pairs of variables below the diagonal; and Pearson’s correlation coefficients above the diagonal. Glioma grade II = pink, grade III = green, Glioblastoma = blue, Metastasis = purple.

The distributions of several of the variables are severely skewed, and many of the bivariate scatterplots reveal nonlinear relationships. As pre-processing of the data to approximate normality and linearity is advantageous in analysis, we used Box-Cox analysis to find transformations X^β^ to achieve this. In all cases we rounded the Box-Cox estimate of the power *β* to a reasonable number. The transformations shown in ***Table 2*** were applied.

**Table 2.** Power β used in Box-Cox transformations.

| $\beta$ | Parameter(s) |
| --- | --- |
| 0.25 | 5, 6, 7 |
| 0.5 | 1 |
| 1.5 | 3 |
| 2.5 | 4 |
| 3.5 | 2 |
| 4.5 | 8 |

The plot matrix of the transformed variables is shown in **Fig. 3**.

**Figure 3.**
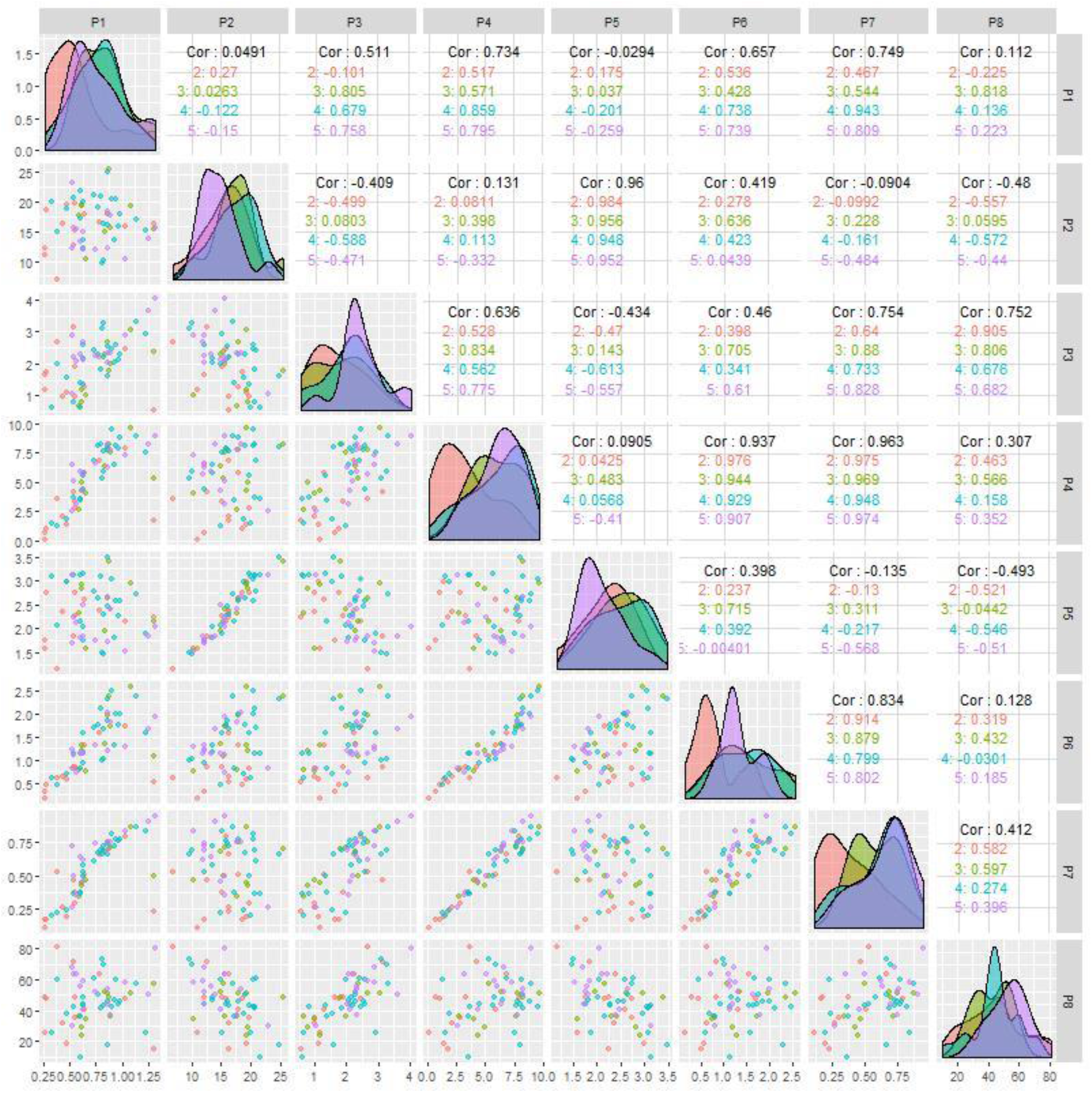
Matrix of plots of transformed data. Density estimates by type are shown on the diagonal; bivariate scatterplots between pairs of variables below the diagonal; and Pearson’s correlation coefficients above the diagonal. Glioma grade II = pink, grade III = green, Glioblastoma = blue, Metastasis = purple.

The distributions are generally closer to normality, and relationships have been linearized in many cases (with a resultant increase in Pearson’s correlation). Pairs of transformed variables exhibiting very high linear correlation (> 0.9) are P2 and P5, P4 and P6, and P4 and P7, showing high correlations between Euclidean parameters and fractal ones (above all, 3D FD and SWI 3D FD). P1-P4, P1-P6 and P3-P6 pairings had moderate correlations. The principal components analysis on the transformed variables gave the percent of variance explained as shown in ***Table 3***. Three principal components explain 94.1% of the variance; four PCs explain 97.6%.

**Table 3.** Principal component analysis of transformed variables.

| Principal component | Percent of variance | Cumulative percent |
| --- | --- | --- |
| 1 | 51.6 | 51.6 |
| 2 | 33.9 | 85.5 |
| 3 | 8.6 | 94.1 |
| 4 | 3.5 | 97.6 |
| 5 | 1.6 | 99.2 |
| 6 | 0.5 | 99.7 |
| 7 | 0.2 | 99.9 |
| 8 | 0.1 | 100.0 |

We investigated the use of the following discrimination methods on the PCs: linear discriminant analysis, quadratic discriminant analysis, K-nearest neighbors, and support vector machines (SVM), based on either three or four principal components. SVMs based on four principal components yielded the best results. The classification of the actual types is shown in ***Table 4***. The overall percent of cases correctly classified by the method was 35/61 = 57%. Glioblastoma had the best classification rate of 88% and grade III glioma the worst at 20%.

**Table 4.** Classification of actual types using support vector machine method on four principal components.

| Actual type | Predicted type |  |  |  | Total | Percent correctly classified |
| --- | --- | --- | --- | --- | --- | --- |
|  | Glioma II | Glioma III | GBM | Metastasis |  |  |
| Glioma II | 7 | 0 | 6 | 0 | 13 | 54% |
| Glioma III | 1 | 2 | 7 | 0 | 10 | 20% |
| Glioblastoma | 2 | 0 | 21 | 1 | 24 | 88% |
| Metastasis | 0 | 0 | 9 | 5 | 14 | 36% |

## 4 Discussion

SWI sequences provide information on venous vasculature, hemorrhage, iron deposits, and very small vascular structures within the central nervous system (including brain tumors) [4, 15–17]. Fractal geometry has been demonstrated to offer appropriate tools to quantify irregular-shaped biological objects, including microvascular and SWI patterns [8], that can be used in automated algorithms for counting microbleeds [18], quantitative susceptibility mapping [19], percentage-wise quantification of intratumoral-susceptibility signals [20], local image variance [21] and other quantitative methodologies [16]. The fractal dimension, the most used parameter in fractal geometry, has been shown as a reliable numerical index to objectively quantify geometrical complexity of microvascular patterns in brain tumors [22]. The fractal dimension is a characteristic of irregularly shaped structures that maintains a constant level of complexity over a limited range of scales [8, 23]. Our previous findings showed that lower FD values of the intratumoral SWI patterns are associated with tumoral microvasculature, while higher values (more “space-filling” and geometrically complex) are associated with microbleeds and necrotic areas [5].

Although not discriminative, the computed parameters quantified in an objective way some intratumoral features, i.e., the SWI patterns of gliomas and brain metastases. This means that the specific fingerprint related to tumors can be quantified by means of radiomics fractal-based parameters. Although an overall percent of 57% cases correctly classified by the method appears to be low, it has to be emphasized that SWI is generally not used for grading and typing in the clinical setting of brain tumors neuroimaging. Nevertheless, SWI’s capacity to visualize intratumoral microstructural features can be linked with quantitative tools, with the aim to characterize the neoplastic architecture and eventually follow it up over time. Advantages include: once the pipeline is standardized, the computed parameters are objective and independent from any operator-dependent variability; the methodology lacks the necessity to augment data when the dataset is small, as it is instead required in AI-based computation of small samples. Limitations are related to the fact that different protocols to acquire SWI features may create a variability in the inputs to be analyzed; moreover, the computation of the parameters is based on the box counting algorithm, which is a valid tool in mathematics and research but not well-known and widespread in the clinical setting. Moreover, the manual segmentation of the ROIs is time-consuming and can be affected by operator-dependent bias. Once again, it is shown that grade III gliomas are the most difficult ones to be differentiate according to their neuroimaging fingerprinting, at least in terms of SWI patterns [5, 10]. Despite such limitations, we believe that the fractal-based computational methodology can become a reliable tool in the neuroradiological armamentarium for the characterization of intrinsic features of brain tumors. Moreover, the present analysis has been applied to an MRI sequence, SWI, which is generally not used for typing and grading of brain tumours. Our results show a proof-of-concept of the applicability of the radiomics analysis to SWI. In future studies, complementary computational analyses will be applied to multimodal imaging for higher diagnostic predictivity.

## 5 Conclusion

A computational pipeline is proposed to provide a standardized diagnostic tool for the medical community. Future studies may include the combination of morphometric Euclidean and fractal parameters, merged with advance image analysis and statistical tools, as well as deep learning methodologies, to objectively characterize brain tumors, at benefit of automatic diagnosis and follow-up of regions of interest in neuroimaging.

## Abbreviations

FD: Fractal Dimension
WHO: World Health Organization
GBM: Glioblastoma
ROI: Region of interest
SWI: Susceptibility-weighted imaging

## Acknowledgments

Prof. Antonio Di Ieva received the 2019 John Mitchell Crouch Fellowship from the Royal Australasian College of Surgeons (RACS), which, along to Macquarie University co-funding, supported the Computational NeuroSurgery (CNS) Lab.

## Ethical approval

All procedures performed in the studies involving human participants were in accordance with the ethical standards of the institutional and/or national research committee and with the 1964 Helsinki Declaration and its later amendments or comparable ethical standards.

## Informed consent

Informed consent was obtained from all individual participants included in the study.

